# The natural selection of metabolism explains curvature in allometric scaling

**DOI:** 10.1101/090191

**Authors:** Lars Witting

## Abstract

I simulate the evolution of metabolism and mass to explain the curvature in the metabolic allometry for placental and marsupial mammals. I assume that the release of inter-specific competition by the extinction of dinosaurs 65 million years ago made it possible for each clade to diversity into a multitude of species across a wide range of niches. The natural selection of metabolism and mass was then fitted to explain the maximum observed body masses over time, as well as the current inter-specific allometry for metabolism. The estimated selection of mass specific metabolism was found to bend the metabolic allometry over time, with the strongest curvature in the placental clade. The rate of exponential increase in mass specific metabolism for placentals was estimated to 9.3 × 10^−9^ (95% CI:7.3 × 10^−9^ − 1.1 × 10^−8^) on the per generation time-scale. This is an order of magnitude larger than the estimate for marsupials, in agreement with an average metabolism that is 30% larger in placentals relative to marsupials of similar size.

## 1 Introduction

The bare existence of an inter-specific allometry between metabolism and mass indicates that the two traits are selected as an energetic balance, where the natural selection of mass is dependent on the selection of metabolism, and/or the selection of metabolism is dependent on the selection of mass. This coordinated natural selection was first considered by Witting (1995, 1997), when he showed that the ecological geometry of the density dependent selection of mass is explaining the exponents of eight inter-specific allometries; including a linear scaling between mass specific metabolism and mass on log-scale, with a −1/4 exponent for two dimensional ecology (2D), and a −1/6 exponent for 3D ecology.

But the metabolic allometry in mammals is not a straight line. It is instead an upward bend curve, where the local exponent for total metabolism is increasing from smaller values around 0:67 to larger values around 0:75 with an increase in the masses of the species that are being compared in the allometric relation (Kolokotrones et al. 2010). A split of mammals into placentals and marsupials illustrates that the curvature is evident in placentals, but apparently not in marsupials that, when two outliers are removed, have an almost perfect Kleiber scaling with an estimated exponent of 0.75±0.01 (MacKay 2011). In order to examine the evolution of these different degrees of curvature in metabolic scaling, I use the selection theory of Witting (2016a,b,c) to simulate the evolution of placental and marsupial mammals over the past 65 million years.

The theory that I use for the simulations was originally proposed as one-sided, in the sense that it described the inuence on metabolism from primary selection of mass (Witting 1995, 1997), but not the inuence on mass from primary selection on metabolism. The former selection was well-suited for the prediction of typical allometries in multicellular animals, but the exponents in unicellular animals extend beyond the predicted range, with the exponent for mass specific metabolism approaching 0:84 in prokaryotes (Makarieva et al. 2005, 2008; DeLong et al. 2010; Witting 2016a).

A two-sided model with primary selection on metabolism and mass was developed by Witting (2016a,b,c). The resource assimilation parameter of the original model was extended into a product, where assimilation by resource handling was multiplied by the pace of the handling process. This product generates gross energy, and by defining the net energy that is available for self-replication as the difference between the assimilated gross energy and the total metabolism of the organism, Witting (2016a) found the mass specific work of handling to be selected as mass specific metabolism. This implies primary selection for an increase in mass specific metabolism; an increase that generates net energy that is a pre-condition for the selection of mass and the associated secondary rescaling of mass specific metabolism with the evolutionary changes in mass (Witting 2016a,b).

Mass specific metabolism and its allometric relation with mass was then found to evolve from primary selection on metabolism and mass (Witting 2016a), with an allometric exponent for mass specific metabolism that declines with a decline in the importance of mass specific metabolism for the generation of the net energy that is selected into mass. This selection predicts a directional evolution of lifeforms from replicating molecules over prokaryote and larger unicells to multicellular animals, with allometric transitions where the exponent for mass specific metabolism is declining from 5/6 (3D) in prokaryotes, over intermediate values in larger unicells, to −1/4 (2D) and −1/6 (3D) in multicellular animals (Witting 2016b).

The prediction of −1/4 and −1/6 exponents for taxonomic clades of multicellular animals is dependent on the assumption that the major component of the body mass variation in the clade originates from primary variation in the handling and/or density of the underlying resources (Witting 2016a). This assumption may hold in many cases at least as an approximation, as it is reasonable to assume that a large fraction of the body mass variation is generated by the adaptation of species to the resources in different niches.

But what happens over evolutionary time when this evolution comes to a hold because the species are becoming well-adapted to their niches? If there is no background evolution in mass specific metabolism we may expect a stationary distribution over time. Yet, as mass specific metabolism is predicted to be selected as the pace of the resource handling that generates net energy for self-replication and the selection of mass (Witting 2016a), we may always expect a certain positive background evolution in mass specific metabolism and mass. And with this paper I aim to analyse whether this background evolution in mass specific metabolism and mass will explain the different degrees of curvature that are observed in the metabolic allometry of placental and marsupial mammals.

## 2 The selection model

To simulate the natural selection of distributions of mammal species that differ in metabolism and mass, I simulate the evolution of placental and marsupial mammals from the extinction of the dinosaurs at the Cretaceous-Palaeogene (K-Pg) boundary 65 million years ago (MA). This involves a two-step model for each clade, with one component that make the clade at the K-Pg boundary diversity into a multitude of lineages in different niches, and another component that is selecting the metabolism and body mass of each lineage in the clade over time.

For simplicity reasons, I assume a single ancestor for each clade at 65 MA, and I let the release of interspecific competition by the extinction of the dinosaurs make it possible for the lineage to diversity into a large clade with a multitude of species across a wide range of niches. Primary selection for an increase in net energy is selecting for the exploitation of the more resource rich niches. And by assuming speciation between populations in different niches, we expect that competitive exclusion by inter-specific interactions will create a distribution of species across niches. The species in the resource rich niches are selected for a higher net energy over time, while species that are excluded into resource space niches may experience a decline in net energy.

### 2.1 The selection of net energy

To describe the selection within the phylogenetic lineage of a species, I let metabolism and mass evolve by the natural selection in Witting (2016a,c). Hence, for unconstrained selection, we have primary selection for an exponential increase

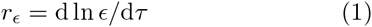

in the average net energy (*ϵ*) on the pre-generation time scale (*τ*) of natural selection. This increase is driven by an exponential increase

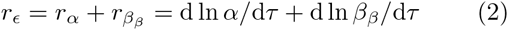

in the two sub-components of resource handling (*α*) and pace 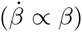, with the latter component being selected as a proxy of mass specific metabolism (*β*), with subscript *β* denoting the pre-mass component of metabolic evolution that occurs independently of the natural selection of mass.

Resource handling (*α*) is given here as a joint function of the actual handling of the resource and of the resource itself. The overall resource handling of a species in a given niche may therefore be selected only towards an upper limit (*α*^**^) that is the maximum handling of the resources in the given niche. A species is fully adapted to its niche when *α* = *α*^**^, and when such a species remains in the niche, it follows that the exponential increase in net energy can continue only by the exponential increase in the metabolic pace of the optimal handling process.

In my modelling I will assume a steady unconstrained natural selection on metabolism across all the species in a clade. This implies that I can expect a rather similar exponential increase in mass specific metabolism 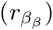 on the per generation time-scale, and I do therefore, for simplicity, assume that all species evolve by the same 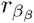 rate. This rate is then estimated separately for placentals and marsupials.

This is unlike the situation for resource handling, where each lineage will evolve towards the niche specific optimum. For this I assume a linear trend

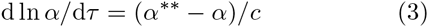

where the rate of exponential change in resource handling is proportional to the evolutionary distance from the optimum, with *c* being a constant. When a species is adapting to a new niche with more or less resource, we expect that the initial change in the net energy that is obtained per handling cycle is large, and that the rate of change will decline as the species approached the optimum.

### 2.2 Life history selection

With the three equations above we can calculate the evolutionary changes in the net energy of the average individual in the population; and it is this change in energy that is driving the natural selection of mass and the associated mass-rescaling of the life history (Witting 2016a,b,c). For multicellular animals like mammals, the increase in net energy is first allocated into replication, where the sustained interference competition that follows from the population growth of self-replication is selecting the energy into an exponential increase in mass (Witting 1997, 2003, 2016c). The associated selection attractor is an evolutionary steady state, where the per generation (*τ*) change in mass (*w*) can be given as

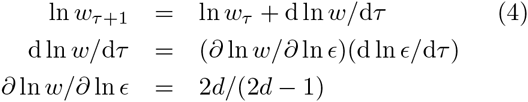

where *d* is the ecological dimensionality of the foraging behaviour, and the log-linear selection relation ∂ ln *w*/∂ ln *ϵ* = 2*d*/(2*d* − 1) is the inverse of the exponent 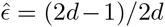 for the deduced allometry between net energy and mass (Witting 2016a), with *d* = 2 for the terrestrial mammals in this study.

The evolution of mass specific metabolism is then given by the primary pre-mass selection (d ln *β*_*β*_/d*τ*) of eqn 2 and the mass-rescaling response of metabolism (d ln *β*_*w*_) to the evolutionary changes in mass (Witting 2016a). Hence,

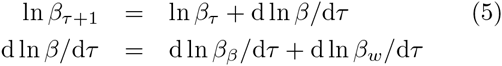

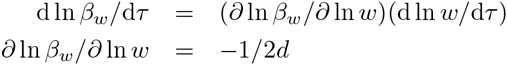

where ∂ ln *β*_*w*_/∂ ln *w* = −1/s*d* is the linear selection relation of the deduced mass-rescaling exponent for mass specific metabolism.

To transfer 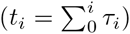 these evolutionary trajectories that are calculated on the per generation (*τ*) time-scale of natural selection into physical time (*t*), we need also to calculate the per generation changes in *τ*. These changes

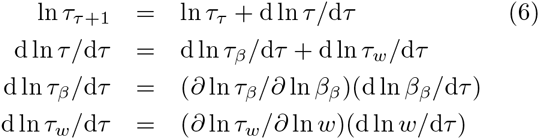

are given by the mass (d ln *τ*_*w*_/d*τ*) and metabolic (d ln *τ*_*β*_/d*τ*) rescaling changes in generation time in response to the evolutionary changes in mass and the pre-mass component of mass specific metabolism, where ∂ ln *τ*_*w*_/∂ ln *w* = 1/2*d* is the selection relation of the deduced mass-rescaling exponent, and ∂ ln *τ*_*β*_/∂ ln *β*_*β*_ = −1 is the rescaling of the generation time in relation to the pre-mass evolution of mass specific metabolism (Witting 2016a).

### 2.3 Mammalian evolution

To simulate the evolution of the inter-specific metabolic allometry for placental and marsupial mammals, I assumed that the common ancestors were adapted to their niche prior to the K-Pg boundary at 65 MA (with *r*_*α*_ = 0 and *α* = *α*^**^). Following an estimate from O’Leary et al. (2013), I assume a common ancestor with a mass of 125 grams at 65 MA for placentals. This mass is about the same as the geometric mean of 146 grams across all placentals today (for the mass data of Pacifici et al. 2013). And with a current average mass of 309 grams for all marsupials, I assume a 300 grams marsupial ancestor at 65 MA.

Then, at 65 MA I created a set of hundred species of 125 gram placentals, and 100 species of 300 gram marsupials, that diversify into niches that differ linearly in ln *α*^**^ from a minimum 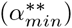 to a maximum 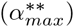. Given the same metabolic pace, it is the potential maximum to resource handling 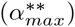 that will allow for the evolution of the largest body masses. The maximum potential resource handling 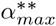 and the *c* parameter of eqn 3 were therefore determined by least-squares fits of the body mass trajectory of 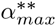 to the global maximum body mass of terrestrial mammals for placentals, and to the maximum for South America for marsupials, with data obtained from Smith et al. (2010).

The potential minimum to resource handling 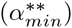 was adjusted to hit a current minimum mass of 2.2 grams for placental and of 7 grams for marsupials. These values corresponds with the minimum masses in the McNab (2008) data that I used for the allometric relationship between metabolism and mass.

The generation times (*τ*_65MA_) of the ancestors at 65MA were adjusted to predict a generation time of 3.3 years for a current placental mammal of 146 grams, and of 2.9 years for a current marsupial mammal of 309 grams. This coincides with the geometric means across all extant placentals and marsupials, as calculated from the body mass and generation time estimates of Pacifici et al. (2013).

I could then for different values of 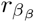 and *β*_65MA_, adjust the 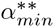, 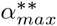, *c* and *τ*_65MA_, parameters as above, and simulate the species distributions over time. The current distributions were transformed by a linear interpolation into a predicted relationship between metabolism and mass, and this allowed me to adjust the 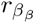 and *β*_65MA_ parameters to obtain the best leastsquares fit between the predicted relationships and the allometric data from McNab (2008), with confidence intervals estimated by bootstraps of the data.

## 3 Results

In the absence of primary selection on metabolism 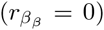, the simulated current relationship between metabolism and mass is Kleiber (1932) scaling, with a log-linear allometry and a 3/4 exponent for the relationship between total metabolism and mass (Fig. 1).

**Figure 1:**
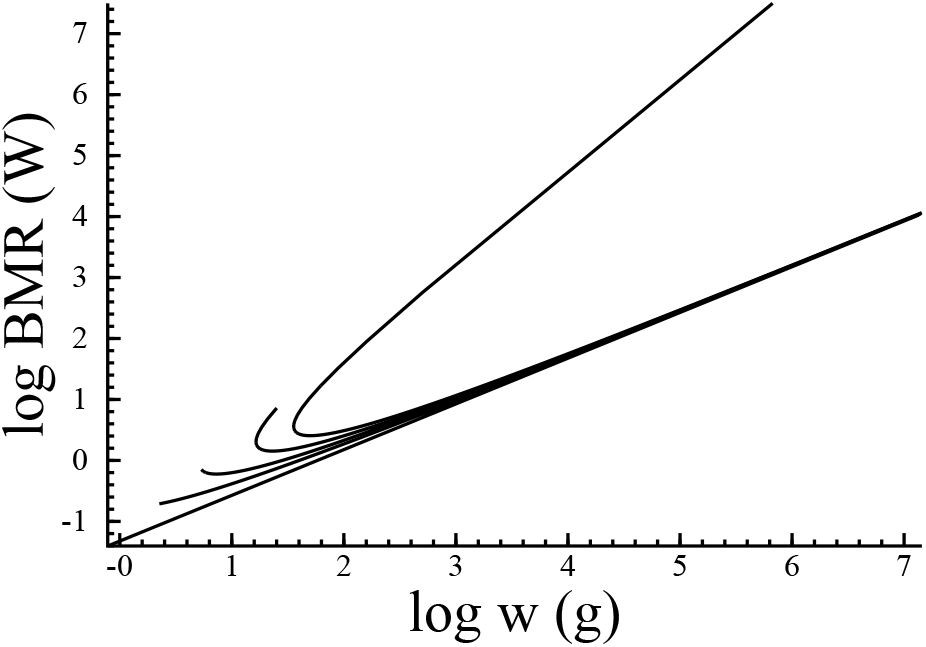
The allometric bend. The predicted un-constrained relationship between the basal metabolic rate (BMR) and body mass (*w*) for a mammalian like clade that has evolved for 65 million years. A straight allometric line with an exponent of 0.75 is expected only in the absence of primary selection on mass specific metabolism 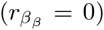. The relationship is instead bend to an increasing degree with an increase in 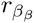, with the illustrated relationships representing a 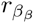 of 0, 1 × 10^−8^, 1.5 × 10^−8^; 2.0 × 10^−8^, and 2.4 × 10^−8^.

With primary selection on metabolism, we find that it is especially the mass specific metabolisms of the smaller species that increase over time. Although all the mammalian lineages in a simulation had the same rate of increase in mass specific metabolism on the per generation time-scale of natural selection, the relative increase is largest in the smaller species in physical time. This is because they, due to their smaller generation times from the mass-rescaling of the body mass evolution just after the K-Pg boundary, evolve over a larger number of generations than the larger species. And as the metabolic increase is selected into mass by eqns 2 and 4, we find that the left-hand side of the allometry is bending upward with respect to both metabolism and mass (Fig. 1).

Initially, for small rates of increase in metabolism, this creates an upward bend in the left-hand side of the allometry. For the larger rates of increase in Fig. 1, it creates the evolution of a separate branch of phylogenetic lineages with large body masses and increasingly high metabolic rates. This second branch has an allometric exponent of 3/2 for total metabolism and of 3/4 for mass specific metabolism.

### 3.1 Placental mammals

The simulated span of body masses for placental mammals over the past 65 million years are shown in the top plot in Fig. 2 for the best estimate of the exponential increase in metabolism. The inter-specific metabolic allometries that follow from the simulation at 50MA, 30MA and 0MA are shown in the bottom plot together with the data for terrestrial placentals from McNab (2008).

**Figure 2:**
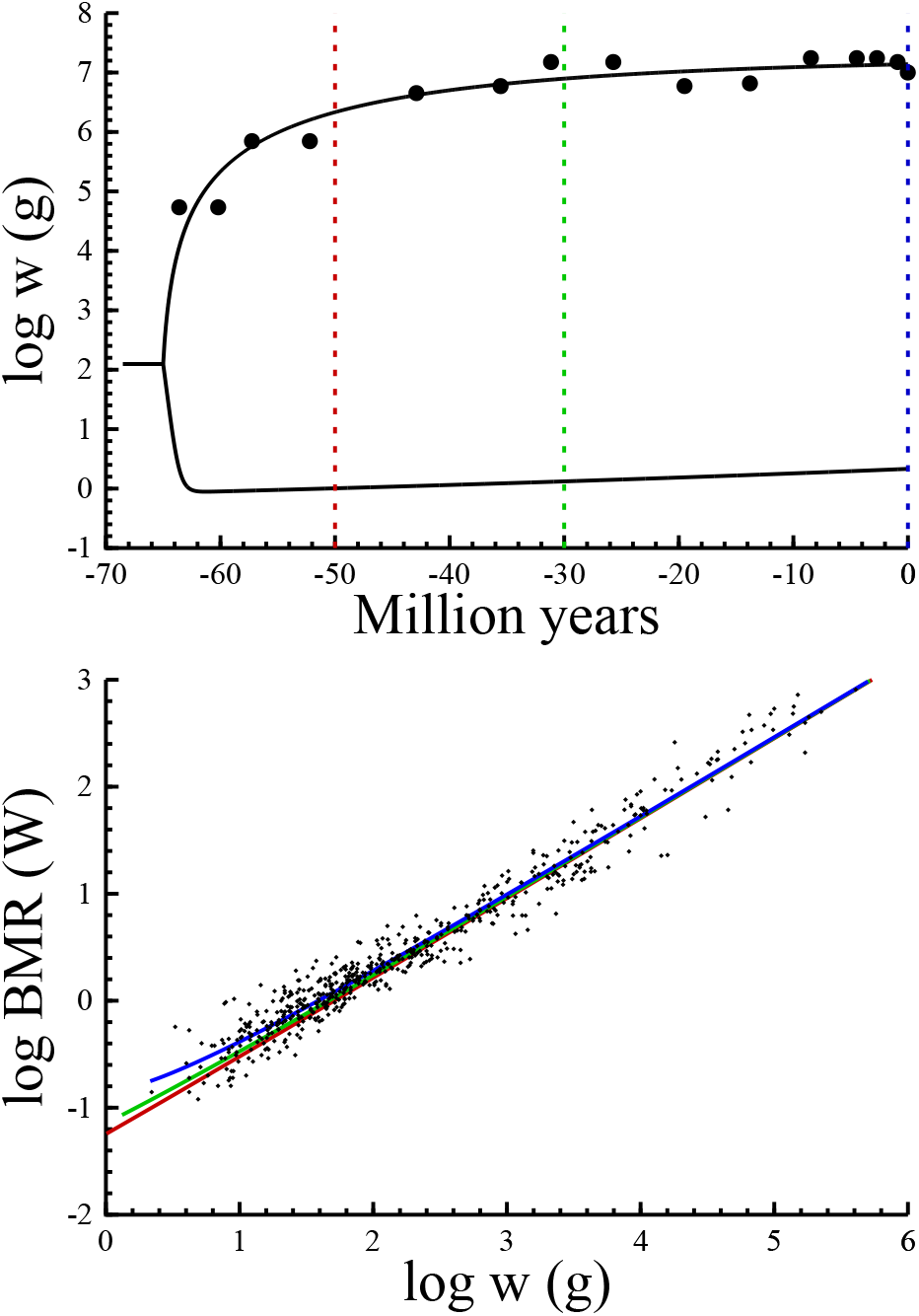
Placental mammals. Top: The span of the simulated body mass (*w*) distribution over time (curves), with dots being the global maximum estimates from Smith et al. (2010). The dashed colour lines mark the time of the simulated allometries in the bottom plot. **Bottom**: The simulated (colour curves) and observed (dots) relationship between the basal metabolic rate (BMR) and body mass (*w*). Red curve: 50 million years ago (MA); Green: 30MA; Blue: 0MA. Data are terrestrial placentals from McNab (2008).

We find that the estimated allometry is bend by time due to the exponential increase in metabolism. Initially at 50MA just after the evolutionary diversification into a multitude of niches, the evolutionary variation in body mass is selected predominantly from variation in the handling of the different resources. This produces a typical Kleiber allometry with almost no bend from primary variation in mass specific metabolism. But the generation of body mass variation from evolution in resource handling comes to a stop over time, while the selected increase in metabolism continues to feed the natural selection of mass. This is affecting especially the smaller species that are selected over a larger number of generations, and this is apparent from the minimum mass in the top plot of Fig. 2. At first, it declines due to the competitive exclusion of lineages into resource sparse niches. But after a while it begins to increase from the selection increase in mass specific metabolism. The result is an inter-specific allometry that becomes more and more bend over time by the primary selection on mass specific metabolism.

The best fit of the current allometry is given by the blue curve in the bottom plot. The upward bend in the metabolism of the smaller placental mammals implies an overall exponent that is smaller than 0:75 should a linear allometry be fitted to the simulated data. The overall linear exponent is 0.72 across the entire range of simulated body masses, and it increases to 0.74 for the upper half of the body mass distribution, and declines to 0.67 for the lower half. The estimated rate of exponential increase in the pre-mass component of mass specific metabolism 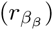 is 9.3 × 10^−19^ (95% CI:7.3 × 10^−9^ − 1.1 × 10^−8^) on the per generation time-scale.

To examine whether the observed bend was caused mainly by the contrast between the largest and smallest placentals, I truncated the data to include the range from 7 g to 32 kg so that it corresponded with the mass range in the marsupial data. This provided a slightly larger point estimate of 1.1 × 10^−8^ that shows that the bend is equally present in the middle rage of the mass data for placentals.

Sensitivity analysis showed that the estimated 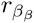 is invariant with respect to deviations in the mass of the ancestor. Yet, the absolute value of 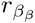 is strongly dependent on the assumed generation time; with an increase in the initial generation time by 20% generating an approximate 20% increase in the estimate.

### 3.2 Marsupial mammals

The simulated span of body masses for marsupial mammals over the past 65 million years are shown in the top plot in Fig. 3 for the best estimate of the exponential increase in metabolism, with the evolution of the inter-specific allometry shown in the bottom plot. The bend in marsupials is clearly smaller than in placentals, with the per generation estimate of the exponential increase in mass specific metabolism being 3.1 × 10^−9^ (95% CI:4.6 × 10^−21^ - 5.4 × 10^−9^)

**Figure 3:**
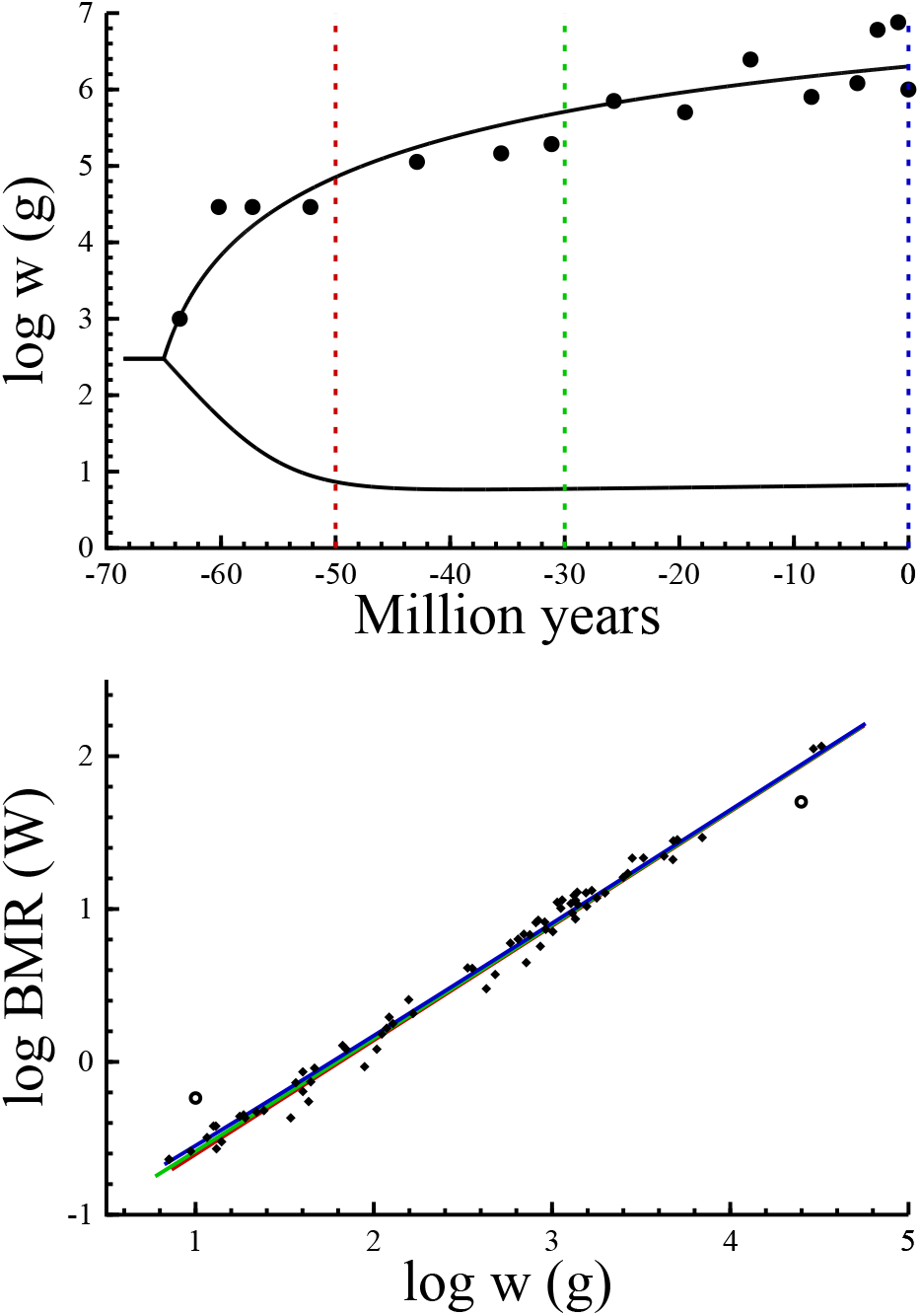
Marsupial mammals. Top: The span of the simulated body mass (*w*) distribution (curves), with dots being the estimates of the maximum mass in South America from Smith et al. (2010). **Bottom:** The simulated (colour curves) and observed (dots) relationship between the basal metabolic rate (BMR) and body mass (*w*). Data from Mc-Nab (2008), with two outlier species (*Tarsipes rostratus* and *Lasiorhinus latifrons*) marked by open circles. See Fig. 2 and text for more details.

This estimate is strongly dependent on the two outlier species that were identified by MacKay (2011); the small honey possum (*Tarsipes rostratus*) with a high metabolism, and the large southern hairy-nosed wombat (*Lasiorhinus latifrons*) with a low metabolism (Fig. 3, bottom). This outlier pattern, with a small species with high metabolism and a large species with low metabolism, is predicted by the primary selection on resource handling, metabolism and mass (Witting 2016c). It occurs in lineages that evolve a small or large body mass at an early stage relative to the other species in the clade. They will then evolve over a larger or smaller number of generations that the main clade and, thus, evolve a higher or lower metabolic rate. When the two outliers are removed, the estimate for 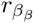 declines by almost an order of magnitude to 6.3 × 10^−10^ (95% CI:4.2 × 10^−22^ - 2.9 × 10^−9^).

The estimate of an average generation time of 2.9 years for a 309 gram marsupial, however, does not match well with an estimate of 3.3 years for an average placental of 146 grams. The average metabolism of a marsupials is only 77% of the metabolism of a similar sized placental (McNab 2008), and this suggests that marsupials should have a longer generation time than placentals. And with a 309 gram placental having an average generation time of 4.0 years, the corresponding allometric estimate for a 309 gram marsupial is 5.1 year. Using this instead of the 2.9 years as a target for marsupials, we obtain a 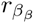 estimate of 1.0 × 10^™9^ (95% CI: 2.9 × 10^−22^) - 5.0 × 10^−9^) for the marsupial clade with the two outlier species removed. This rate is approximately an order of magnitude smaller than the estimate for placentals.

## 4 Discussion

Kolokotrones et al. (2010) found that the relationship between metabolism and mass in mammals is not a straight allometric line, but an upward bend curve that is better described by a second order polynomial. When a straight line was fitted to the larger body masses they observed exponents around 0.75, but the exponent was closer to 0.67 when a straight line was fitted to the smaller masses.

With this paper I have shown that this curvature is explained by primary selection for an exponential increase in the pre-mass component of mass specific metabolism. This evidence for a selection increase in mass specific metabolism is in agreement with other evidence, *i*) where the transitions in the allometric exponents from prokaryotes over larger unicells to multicellular animals are explained from primary selection on metabolism and mass (Witting 2016a), *ii*) where life-forms from virus over prokaryotes and larger unicells to multicellular animals are explained from primary selection on metabolism and mass (Witting 2016b), and *iii*) where the evolutionary bending of mammalian body mass trajectories over millions of years are explained from primary selection on mass specific metabolism (Witting 2016c).

Taken together the evidence suggests that natural selection is generating an intrinsic and exponential acceleration in the pace of the biochemical and ecological processes of mobile organisms; an increase that is selected by an increase in the net energy that the organisms has available for self-replication. This conclusion for a sustained positive selection of mass specific metabolism is in agreement also with an invariance between mass specific metabolism and mass from prokaryotes to mammals (Makarieva et al. 2005, 2008; Kiørboe and Hirst 2014). This invariance indicates that the macro evolution of major taxonomic groups proceed along an upper limit of mass specific metabolism (Witting 2016b), a results that is supported also by a log-linear increase in the maximum mass of mobile organisms over the last 3.5 billion years of evolution (Witting 2016c).

The maximum rates of mass specific metabolism are rather similar across the major taxonomic groups (Makarieva et al. 2005, 2008). Yet, the majority of animals are not selected to the limit. The most important reason for this is probably mass-rescaling (Witting 2016a), where mass specific metabolism is selected to decline with a selected increase in the mass of many species when a phylogenetic lineage diversifies into a multitude of more resource rich niches. Following this diversification in mammals at the K-Pg boundary, and the associated evolution of larger masses and smaller metabolic rates, I simulated the background selection of mass specific metabolism in placentals and marsupials to estimate the degree of bending of the inter-specific allometry over time.

In agreement with MacKay (2011), the bend was found to be more apparent in placentals than marsupials. This difference was estimated to reflect a per generation rate of increase in mass specific metabolism that is about a one order of magnitude larger in placentals. Hence, from the differences in the curvature of the metabolic allometry, we conclude that placentals have evolved a higher metabolism than marsupials; in agreement with an average metabolism that is 30% larger in placentals relative to marsupials of similar size (McNab 2008).

